## Supplementary Information for "Being noisy in a crowd: differential selective pressure on gene expression noise in model gene regulatory networks"

This document contains all supplementary figures and information referenced in the main text. The simulation results data and the code necessary to reproduce all figures is available at `https://doi.org/10.5281/zenodo.6939845`, together with the the code necessary to generate all raw simulation files.

**4 Robustness of results in different topology structures** **23**

**5 Filtered datasets** **25**

### 1 Gene regulatory network model and evolutionary model

#### 1.1 Parameters

The parameter values used for the gene regulatory network model and evolutionary simulations and their descriptions are shown in Table S1.

**Table S1. Parameters used in the simulations.**

| Parameter | Symbol | Value | Description |
|---|---|---|---|
| Number of nodes in the network | $n$ | 40 | Number of genes in the gene regulatory network |
| Network density | $d$ | 0.05 | Proportion of potential connections in the network |
| Regulatory matrix | $W = (w_{ij})_{1\leq i\leq n,\, 1\leq j\leq n}$ | see Supp. Data | Regulatory relationships in the gene regulatory network |
| Intrinsic noise | $\{\eta_i^{\text{int}}\}_{1\leq i\leq n}$ | 100 | Gene-specific noise of each gene |
| Basal expression levels | $\{S_i^{basal}\}_{1\leq i\leq n}$ | $\{20, ..., 20\}$ | Constitutive expression level |
| Number of timesteps for genotype realization | $\text{T}_r$ | 50 | Number of timesteps the expression levels are updated |
| Minimal expression level | $s_{min}$ | 0 | Minimal expression level |
| Maximal expression level | $s_{max}$ | 100 | Minimal expression level |
| Number of timesteps to check oscillatory dynamics | $\tau$ | 10 | Time window to apply oscillation criterion |
| Maximal allowed fluctuation in gene expression levels | $\epsilon$ | $1e^{-06}$ | Criterion to check oscillatory dynamics |
| Population size | $N$ | 1,000 | Number of individuals in a population |
| Number of generations | $T$ | 10,000 | Length of the evolutionary simulation in generations |
| Optimal expression levels (for network establishment) | $\{s_i^{opt}\}_{1\leq i\leq n}$ | $\{50, ..., 50\}$ | Expression levels that correspond to maximum fitness |
| Mutation rate (regulatory interactions) | $\mu_w$ | 0.05 | Mutation probability of a regulatory interaction, per interaction, per repl. event |
| Mutation value mean (regulatory interactions) | $m_w$ | 0 | Mean of normal distribution from which mutation values are drawn |
| Mutation value variance (regulatory interactions) | $v_w$ | 2 | Variance of normal distribution from which mutation values are drawn |
| Selective pressures | $\{\rho_i\}_{1\leq i\leq n}$ | $\{1, ..., 1\}$ | Contribution of expression level to fitness |
| Mutation rate (intrinsic noise) | $\mu_\eta$ | 0.01 | Mutation probability of intrinsic noise, per gene, per repl. event |
| Mutation value mean (intrinsic noise) | $m_\eta$ | 100 | Mean of normal distribution from which mutation values are drawn |
| Mutation value variance (intrinsic noise) | $v_\eta$ | 40 | Variance of normal distribution from which mutation values are drawn |
| Recombination rate | $r$ | 0.05 | Probability of offspring entering the recombination process |

#### 1.2 Robustness of network realization

In the simulations performed in this study the gene regulatory network model was realized into the phenotype by synchronously updating the expression levels of all genes in every time step. To test whether the time of updating changes the steady state expression levels, we compared steady state expression levels of 200 samples of 20-gene random networks realized without noise synchronously and asynchronously. Synchronous updating was performed by updating the expression level of all genes in the network at every time step for $T_r$ time steps ($T_r = 50$). Asynchronous updating was performed by randomly choosing a gene in every time step and updating only its expression levels, for $n \times T_r = 1000$ time steps. We realized each network topology 1000 times synchronously and asynchronously and measured the Hamming distance between the expression level vectors in the last time step. Two expression level values were deemed identical if their difference was less than 0.001. In 92% of cases the synchronous and asynchronous realizations of the same network configuration had a Hamming distance of 0 (Fig S1A). The mean expression level, expression variance, CV, noise and Fano factor were highly correlated between the synchronous and asynchronous realizations (Fig S1B-F). Examples of expression level dynamics of deterministic and stochastic realizations with synchronous and asynchronous updating schemes in one network are shown in Fig S2.

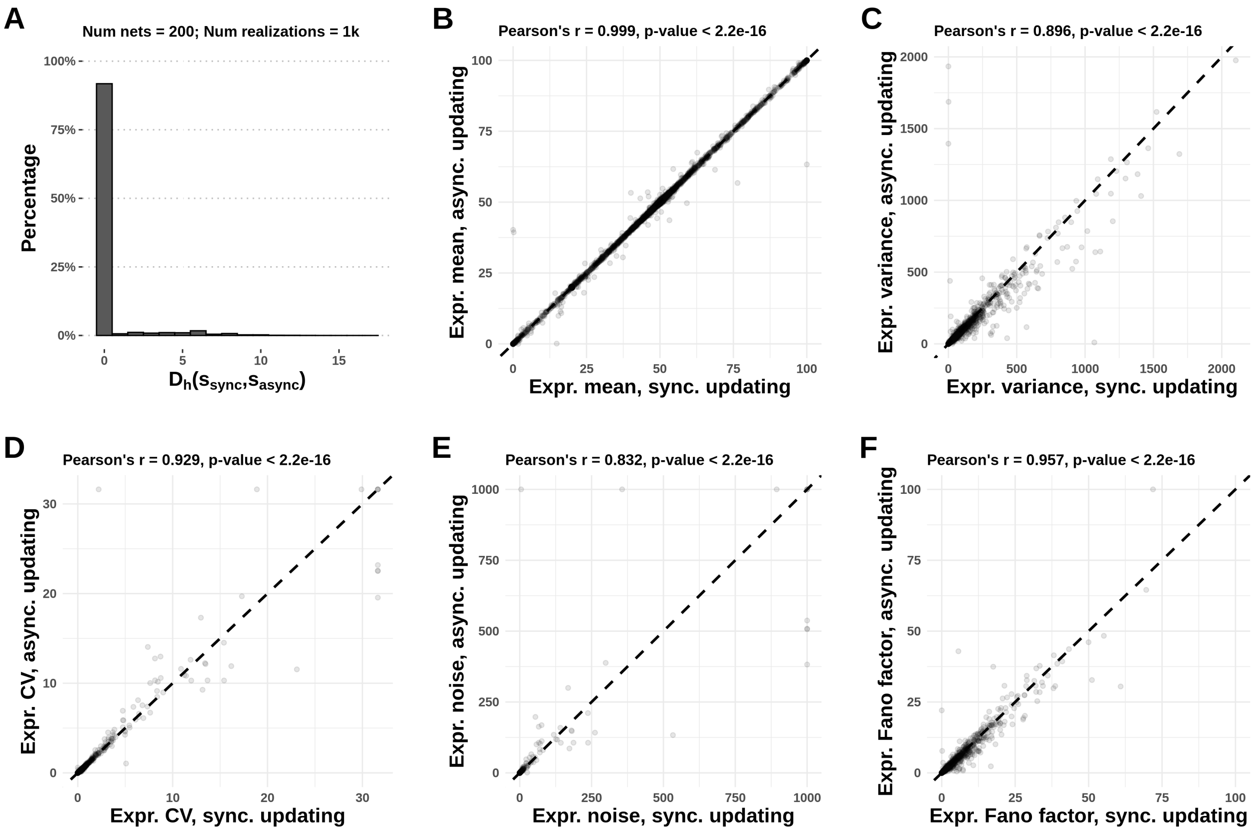

**Fig S1. Network realization is robust to synchronous or asynchronous expression level updating mode during network realization.** **A** - Hamming distance between the steady states of synchronously and asynchronously realized networks. Expression level values between synchronous and asynchronous realizations were deemed identical if their difference was less than 0.001. **B** - Mean expression level of populations of synchronously and asynchronously realized networks. **C-F** - Expression level variance, CV, noise and Fano factor of genes from populations of synchronously and asynchronously realized networks.

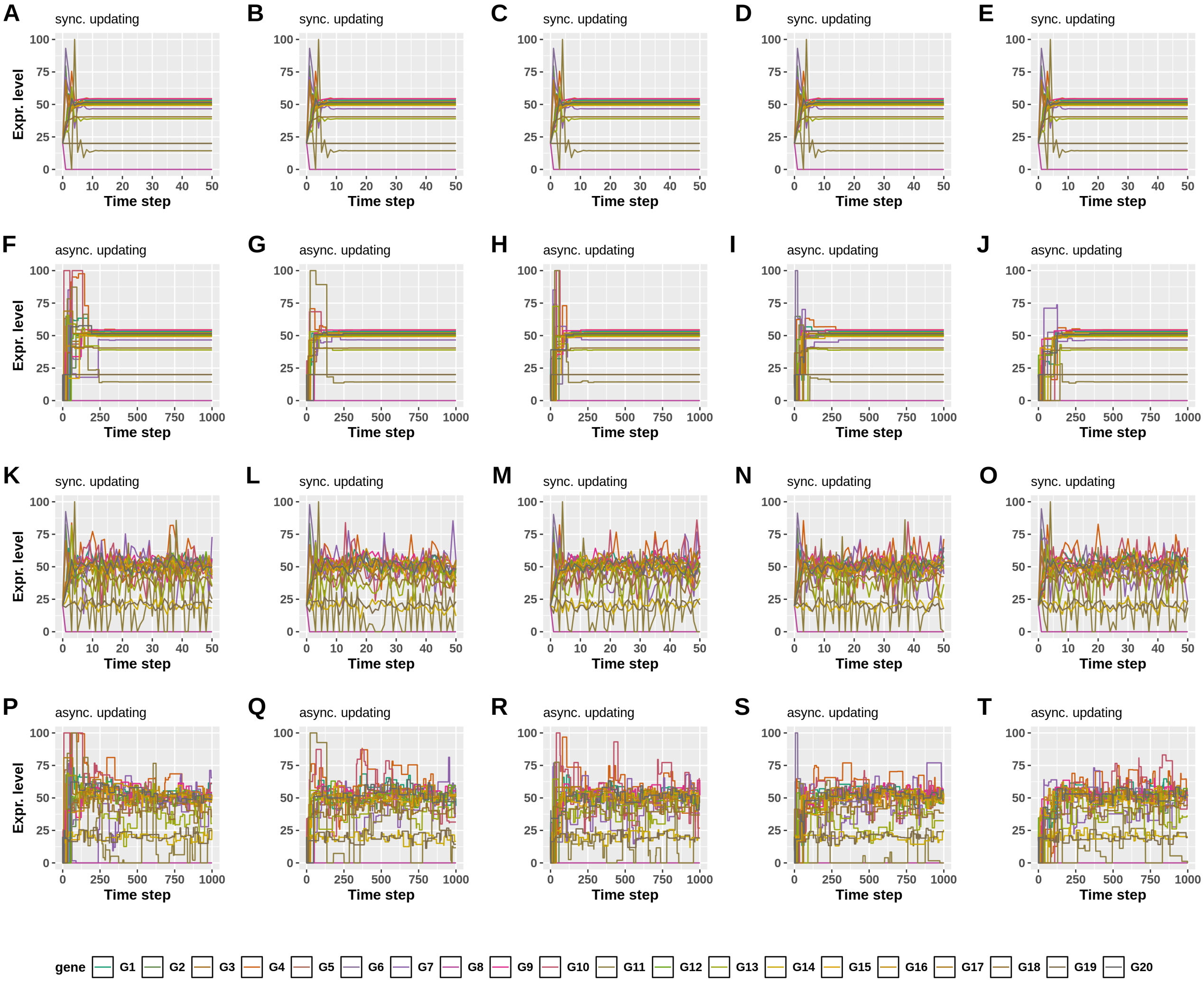

**Fig S2. Examples of expression level dynamics in realizations of the same network with different expression level updating modes and noise levels. A-E** Five non-noisy realizations with synchronous expression level updating. Since there is no random component in the realization, there are no differences between the realizations. **F-J** Five non-noisy realizations with asynchronous expression level updating. The expression level of a randomly chosen gene is updated in each timestep. Consequently, even though there is no intrinsic expression noise, the dynamics differ between the five realizations, but they reach the same steady state as in the synchronously updated realizations. **K-O** Five noisy realizations with synchronous expression level updating. The mean of the gene expression levels equals the steady state expression levels of the realizations without noise. **P-T** Five noisy realizations with asynchronous expression level updating.

#### 1.3 Population size

We tested the effect of the population size on selective pressure by simulating the evolution of a dataset of 500 network topologies with different population sizes. We find that increasing the population size increases the selective pressure acting on constituent genes (Fig S3). A population size of 1000 was chosen for the main simulations in this study.

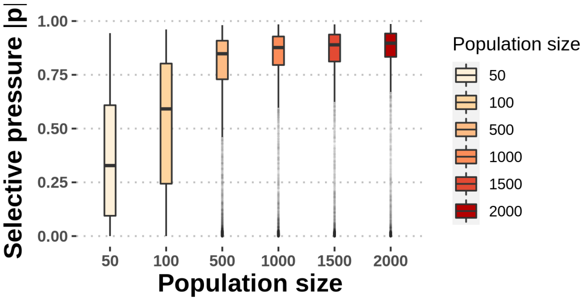

**Fig S3. Increasing the population size increases the selective pressure on genes under stabilizing selection on gene expression level.** Dataset consists of 20,000 genes from 500 random 40-gene network topologies.

#### 1.4 Stability of mean expression level

We imposed stabilizing selection on gene expression levels and observed a repeatable pattern of reduction of gene expression variance. The mean expression level was stable (Fig S4A) throughout evolution, meaning that the adapted populations had a higher fitness due to a reduction of gene expression variance, not changes in expression mean.

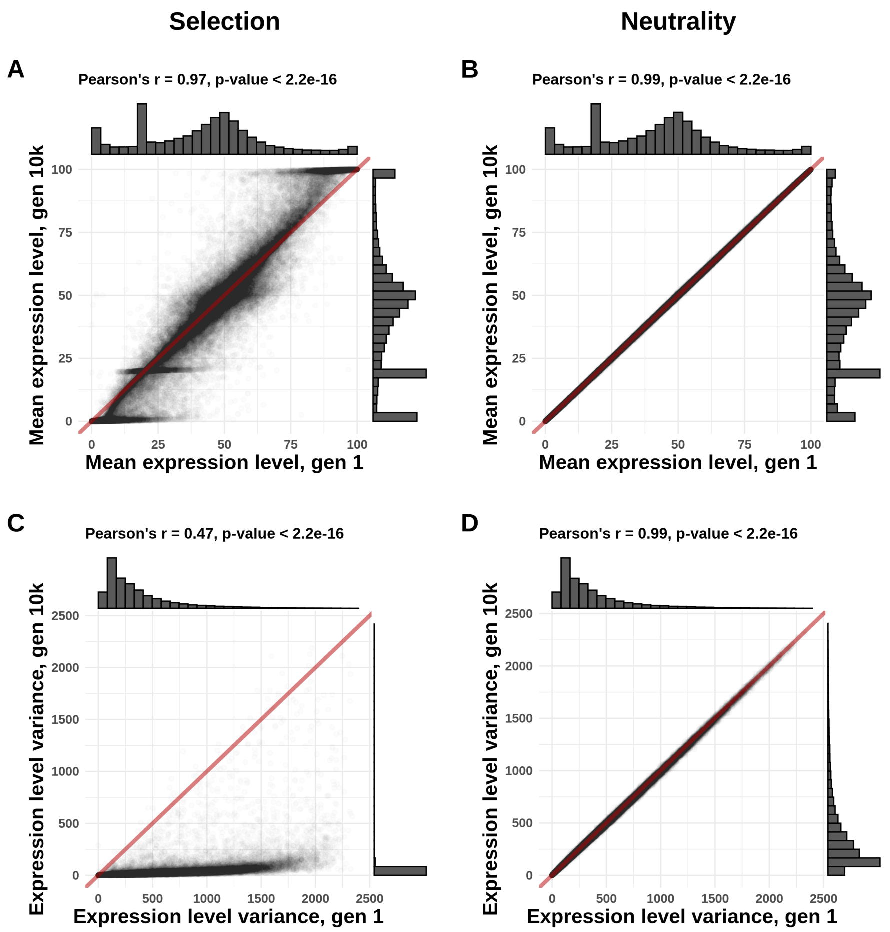

**Fig S4. Mean expression level does not change after noise evolution under stabilizing selection on gene expression levels. A, B** - Mean expression level in the first and last generation of populations evolved under selection (A) and neutrality (B). **C, D** - Expression variance in the first and last generation of populations evolved under selection (C) and neutrality (D). Red lines indicate lines with a slope of 1.

#### 2 Network centrality metrics

To measure the centrality of nodes in the gene networks, we computed 19 node-level centrality measures. These centrality measures are: degree, indegree, outdegree, closeness, betweenness, eigenvector centrality, node strength, instrength, outstrength, hub score, authority including weights, authority excluding weights, absolute node strength, absolute instrength, and absolute outstrength, flow betweenness, load centrality, information centrality, and stress centrality. These measures were heavily intercorrelated and correlated with the expression noise metrics - expression noise, change of expression noise after selection, and selective pressure (Fig S5). We also computed 12 graph-level centrality measures to study the effects of the global topology on the average selective pressure. These measures are: diameter, mean path distance, degree assortativity, degree centralization, indegree centralization, outdegree centalization, closeness centralization, betweenness centralization, average degree, average indegree, and average outdegree. The global network metrics were intercorrelated, as well (Fig S6). In the study of the effects of network centrality on evolvability of gene-specific expression noise we focused on instrength and outstrength as node-level centrality measures, and summarized the 12 graph-level measures into two synthetic independent variables using principal component analysis.

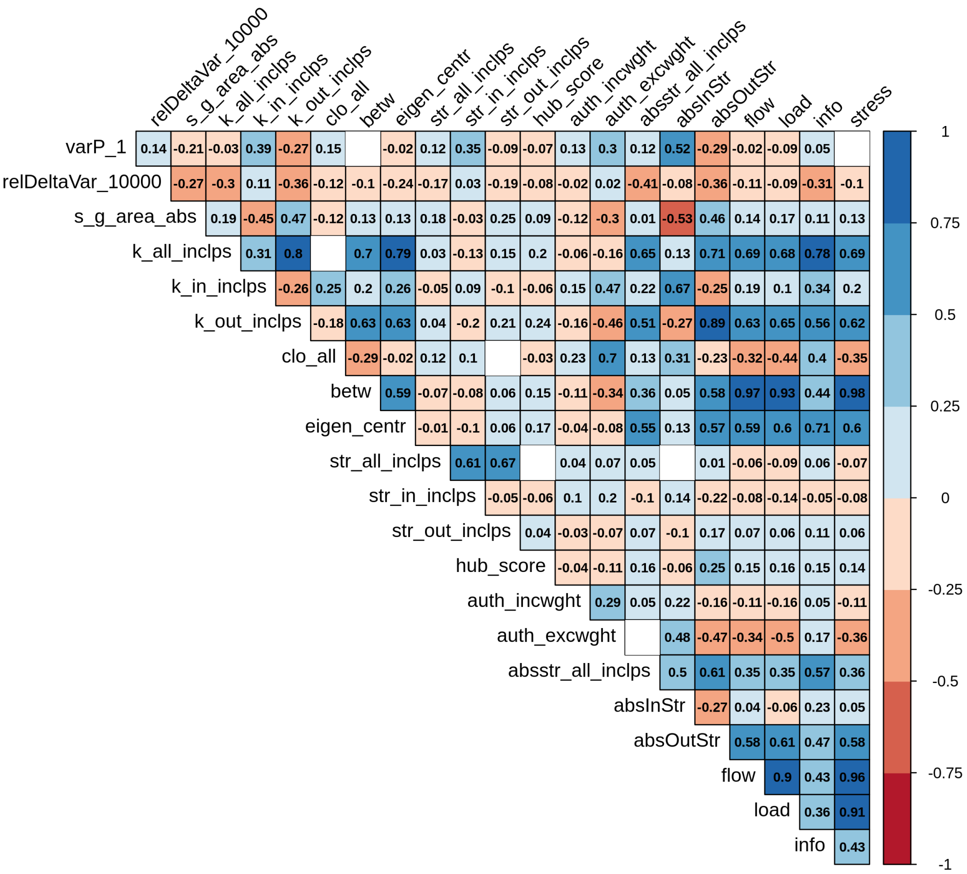

**Fig S5. Correlation matrix of node-level network centrality metrics and expression noise metrics.** Spearman's rank correlation coefficients shown in the cells. Empty cells indicate a non-significant p-value (p-value > 0.05). Abbreviations: varP_1 - expression variance in the first generation; relDeltaVar_10000 - relative change of expression variance between the first and generation 10,000; s_g_area_abs - selective pressure on each node; k_all_inclps - degree; k_in_inclps - indegree; k_out_inclps - outdegree; clo_all - closeness; betw - betweenness; eigen_centr - eigenvector centrality; str_all_inclps - node strength; str_in_inclps - instrength; str_out_inclps - outstrength; hub_score - hub score; auth_incwght - authority including weights; auth_excwght - authority excluding weights; absstr_all_inclps - absolute strength; absInStr - absolute instrength; absOutStr - absolute outstrength; flow - flow betweenness; load - load centrality; info - information centrality; stress - stress centrality. Dataset consists of 148,886 genes from 2,000 random network topologes.

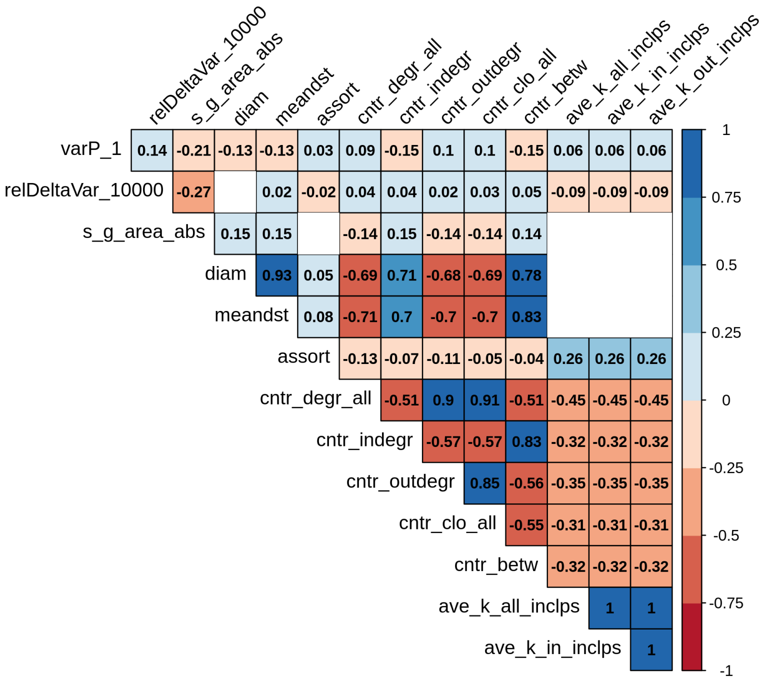

**Fig S6. Correlation matrix of graph-level network centrality metrics and expression noise metrics.** Spearman's rank correlation coefficients shown in the cells. Empty cells indicate a non-significant p-value (p-value $> 0.05$). Abbreviations: varP_1 - expression variance in the first generation; relDeltaVar_10000 - relative change of expression variance between the first and generation 10,000; s_g_area_abs - selective pressure on each node; diam - diameter; meandst - mean path distance; assort - degree assortativity; cntr_degr_all - degree centralization; cntr_indegr - indegree centralization; cntr_outdegr - outdegree centralization; cntr_clo_all - closeness centralization; cntr_betw - betweenness centralization; ave_k_all_inclps - average degree; ave_k_in_inclps - average indegree; ave_k_out_inclps - average outdegree. Dataset consists of 148,886 genes from 2,000 random network topologes..

#### 2.1 Colinearity between instrength and outstrength

The predictor variables used in statistical modelling in the main results, node instrength and outstrength, were correlated (Spearman's $\rho$ = -0.17, p-value $< 2.2 \times 10^{-16}$, Fig S7A-B). This correlation is due to the distributions of in and out nodes being non independent: the more in-connections has, the less out-connections. We also observed that a part of the residuals non-normality may be due to points with a value of zero in one of the two predictor variables. As a control, we rerun the entire analysis on two additional filtered datasets. In the first one, we kept only genes with zero values of either instrength or outstrength, *i.e.* this dataset consisted of only regulators and target genes. Instrength and outstrength were more correlated in the first filtered dataset (Spearman's $\rho$ = -0.86, p-value $< 2.2 \times 10^{-16}$, Fig S7C) than in the unfiltered dataset. In the second filtered dataset, we removed all genes that had a zero value of either instrength or outstrength, *i.e.* this dataset consisted of genes that are both regulators and regulated. Instrength and outstrength were less correlated in the second filtered dataset (Spearman's $\rho$ = -0.03, p-value $< 2.2 \times 10^{-16}$, Fig S7C) than in the unfiltered dataset, and this filtering somewhat reduced the heteroskedasticity of the Pearson's residuals in the statistical models. The same pattern of effects and significance of instrength and outstrength was observed in the filtered datasets as in the main dataset, indicating that our conclusions are robust to the heteroskedasticity of Pearson's residuals and collinearity between the explanatory variables. The results of all statistical models are summarized in Table S6 in Section 5.

#### 2.2 PCA of global network metrics

To investigate the effects of the intercorrelated graph-level network centrality metrics on noise propagation and noise evolution, we performed a principal component analysis (PCA) to construct independent summary variables representing graph-level network centrality metrics. The first two dimensions of the PCA expressed 85.4% of the total data inertia (Fig S8A), so we chose the first two principal components (PCs) as synthetic explanatory variables in linear mixed-effects models in the main results. The loadings of the first two PCs are shown in Fig S8B. The loading of the first synthetic variable (PC1) is dominated by negative loadings of diameter and mean path distance, and the centralization measures, namely positive loadings of outdegree and closeness centralization and negative loadings of indegree and betweenness centralization. The loading of the second synthetic variable (PC2) is dominated by the negative loading of the average degree, average indegree and average outdegree measures. For easier interpretation, the sign of the PCs has been switched in the statistical modelling shown in the main text.

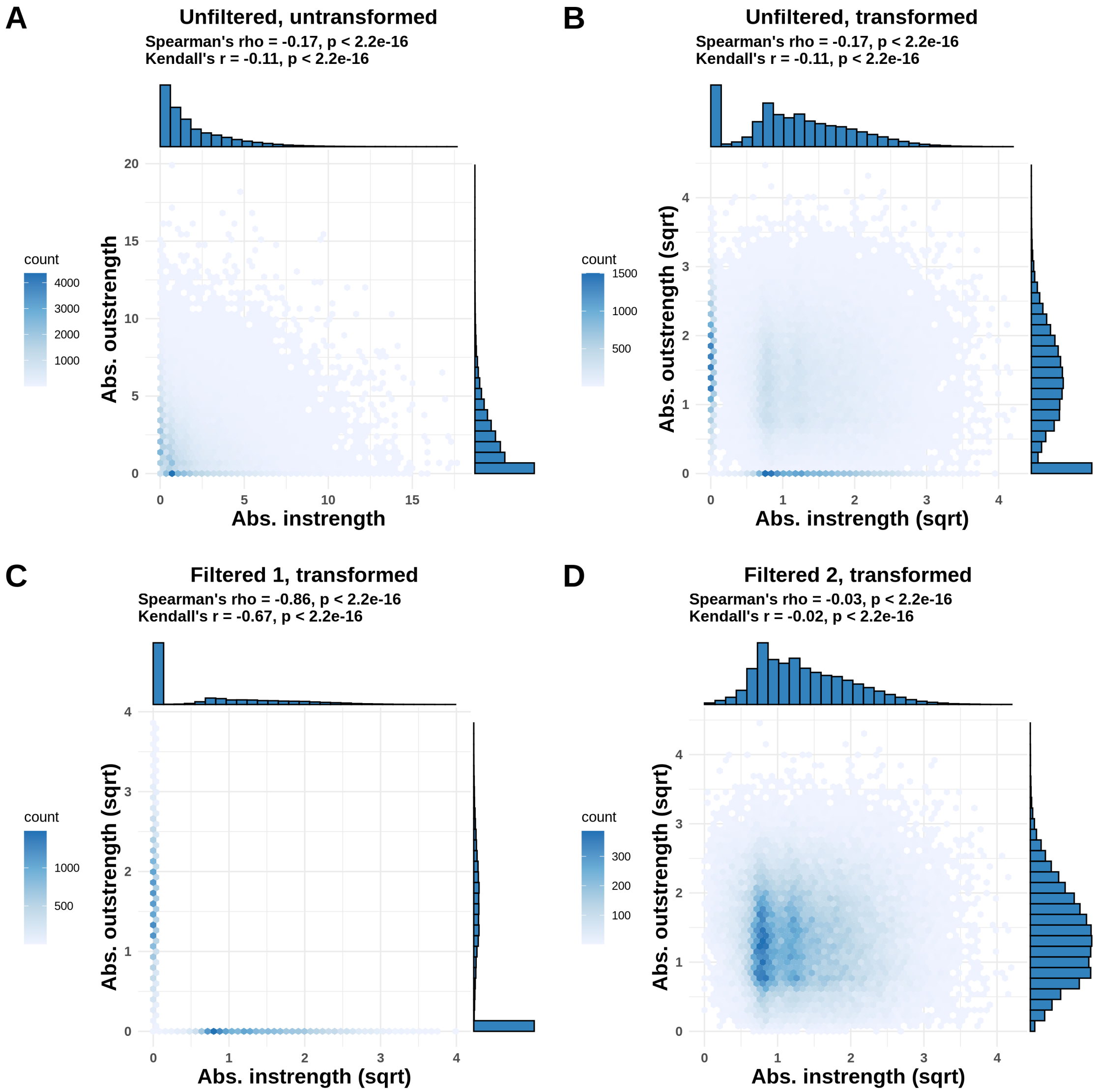

**Fig S7. Correlations between node instrength and outstrength in unfiltered and filtered datasets.** **A** - Correlation between instrength and outstrength in the unfiltered dataset. Dataset consists of 148,886 genes from 2,000 random network topologies. **B** - Correlation between square-root transformed instrength and outstrength in the unfiltered dataset. **C** - Correlation between square-root transformed instrength and outstrength in the filtered dataset. Dataset consists of 43,214 genes from 2,000 random network topologies. **D** - Correlation between square-root transformed instrength and outstrength in the filtered dataset. The dataset consists of 105,672 genes from 2,000 random network topologies.

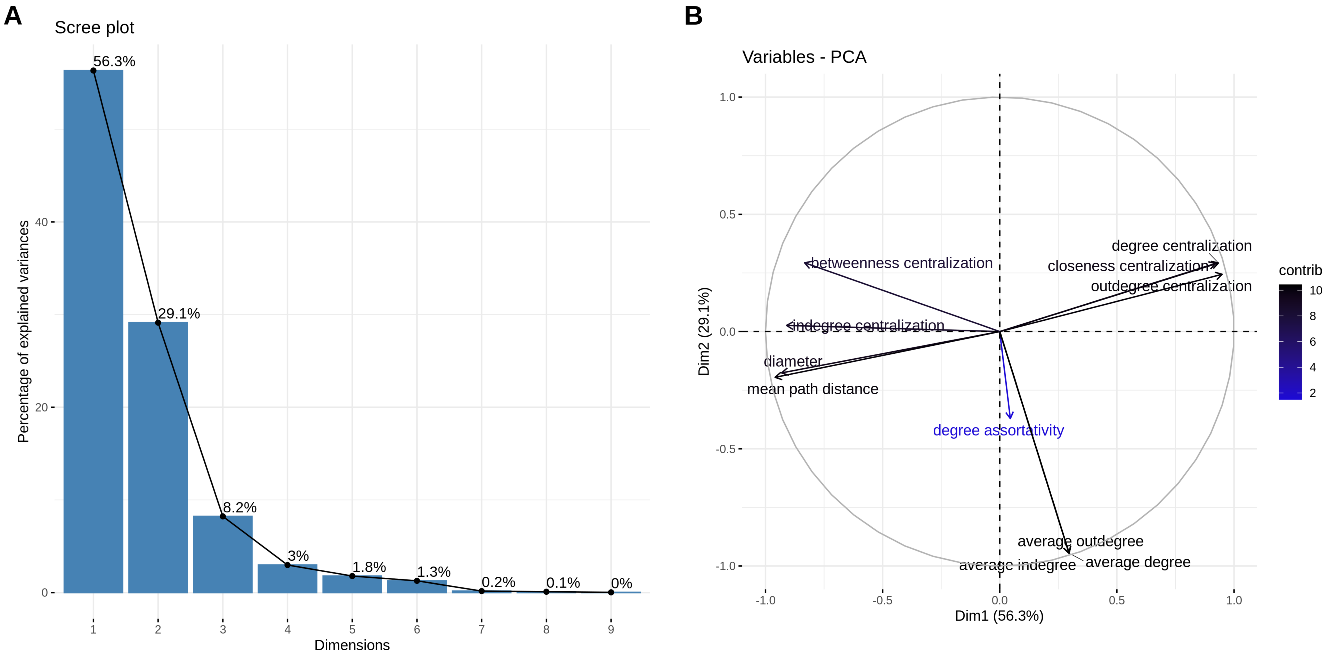

**Fig S8. Principal component analysis of the graph-level network centrality metrics. A** - Scree plot depicting the percentage of total variance explained by each principal component. The first two principal components express 85.4% of the total inertia. **B** - Correlation circle showing the loadings of the first two principal components.

### 3 Diagnostics of statistical models

#### 3.1 GLMM: Noise propagation

To investigate whether noise propagation is captured by our gene regulatory network model, we fitted a linear mixed-effects model with the following formula:

$$y = X\beta + Zu + \epsilon, \tag{1}$$

where the outcome variable, $y$, is a column vector of the expression variance of each node; $X$ is a matrix of two explanatory fixed-effects variables, node instrength and node outstrength, $\beta$ is a column vector of the two fixed-effects coefficients; $Z$ is the column vector for the design of the random effect variable, network topology sample, and the number of groups equivalent to the number of network topology samples; $u$ is a column vector with the random-effects coefficient for each group (network topology sample); $\epsilon$ is a column vector with the residuals. When fitting a model with the assumption of constant variance, the Pearson's residuals were heteroskedastic (Fig S9A). We fitted models with different variance structures and based on Akaike's Information Criterion chose the model with the exponential function of the node instrength as the variance structure. Pearson's residuals of the chosen and all other fitted models are shown in Fig S9B-E. Changing the variance structure did not change the significance or the effect of the fixed variables (Table S2). The variance inflation factor (VIF), a measure of collinearity of explanatory variables, was 1.08. A VIF value lower than 3 indicates that the statistical significance of the inferred effects is reliable in spite of collinearity (1).

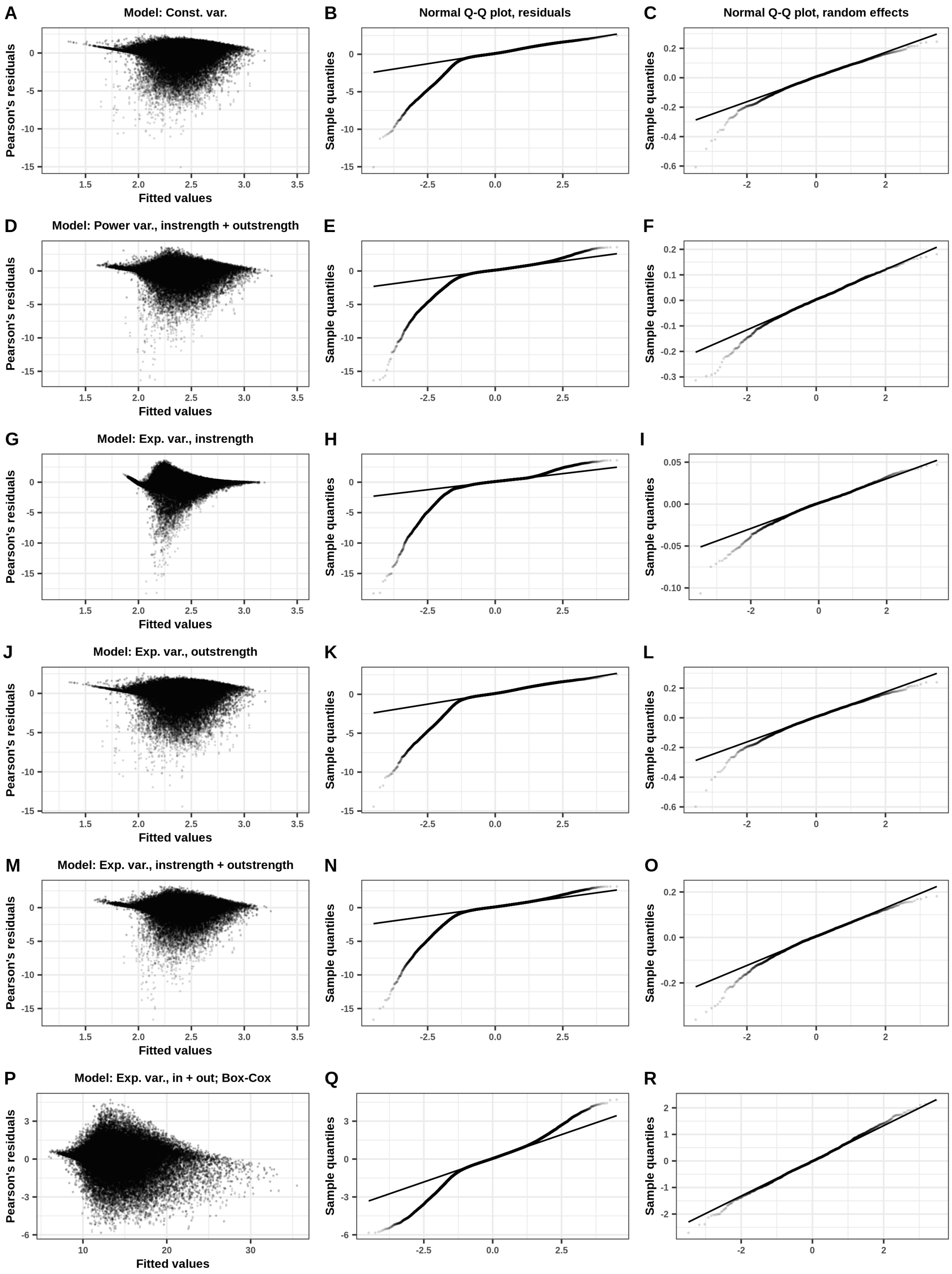

**Fig S9. Diagnostics of linear mixed-effects models with expression variance as a response variable and different variance structures.**

**Fig S9.** Continued. **A-C** Plot of Pearson's residuals vs. fitted values (A), Q-Q plot of Pearson's residuals (B), Q-Q plot of random effects (C) of a model with no variance structure. **D-F** Pearson's residuals vs. fitted values (D), Q-Q plot of standardized Pearson's residuals (E), Q-Q plot of random effects (C) of a model with a variance structure modelled as a power function of instrength and outstrength. **G-I** Pearson's residuals vs. fitted values (G), Q-Q plot of standardized Pearson's residuals (H), Q-Q plot of random effects (I) of a model with a variance structure modelled as an exponential function of instrength. **J-L** Pearson's residuals vs. fitted values (J), Q-Q plot of standardized Pearson's residuals (K), Q-Q plot of random effects (L) of a model with a variance structure modelled as an exponential function of outstrength. **M-O** Pearson's residuals vs. fitted values (M), Q-Q plot of standardized Pearson's residuals (N), Q-Q plot of random effects (O) of a model with a variance structure modelled as an exponential function of instrength and outstrength. **P-R** Pearson's residuals vs. fitted values (P), Q-Q plot of standardized Pearson's residuals (Q), Q-Q plot of random effects (R) of a model with a variance structure modelled as an exponential function of instrength and outstrength and with the explanatory variables transformed with the Box-Cox transform.

**Table S2. Different variance structures do not affect the sign of the effect and significance in linear mixed-effects models with expression variance as a response variable.** The results of models with different variance structures are shown in the table. The effect size differs by a small margin, but the sign and significance remain the same regardless of variance structure. The model with the variance structure as an exponential function of instrength had the lowest Akaike's Information Criterion and was chosen as the best model. Abbreviations: const. var. - constant variance; power var., in + out; variance as a power function of instrength; exp. var., in - variance as an exponential function of instrength; exp. var., out - variance as an exponential function of outstrength; exp. var., in + out - variance as an exponential function of instrength and outstrength; absInStrT_sqrt - absolute instrength, square-root transformed; absOutStrT_sqrt - absolute outstrength, square-root transformed.

| Model | Predictors | Value | Std.Error | p.value | p.significant | AIC |
|---|---|---|---|---|---|---|
| const. var. | (Intercept) | 2.12252974 | 0.0037040699 | 0.000000e+00 | ✔ Yes | 173025.99 |
| const. var. | absInStrT_sqrt | 0.23326569 | 0.0014753454 | 0.000000e+00 | ✔ Yes | 173025.99 |
| const. var. | absOutStrT_sqrt | -0.07906913 | 0.0014530464 | 0.000000e+00 | ✔ Yes | 173025.99 |
| power var., in + out | (Intercept) | 2.03654963 | 0.0024398194 | 0.000000e+00 | ✔ Yes | 144787.06 |
| power var., in + out | absInStrT_sqrt | 0.27234172 | 0.0012837816 | 0.000000e+00 | ✔ Yes | 144787.06 |
| power var., in + out | absOutStrT_sqrt | -0.04355256 | 0.0011948879 | 1.415734e-289 | ✔ Yes | 144787.06 |
| ★ exp. var., in | (Intercept) | 2.01421296 | 0.0014226309 | 0.000000e+00 | ✔ Yes | 96143.99 |
| ★ exp. var., in | absInStrT_sqrt | 0.27887716 | 0.0011158593 | 0.000000e+00 | ✔ Yes | 96143.99 |
| ★ exp. var., in | absOutStrT_sqrt | -0.02214644 | 0.0007327439 | 4.731613e-200 | ✔ Yes | 96143.99 |
| exp. var., out | (Intercept) | 2.11543714 | 0.0036669065 | 0.000000e+00 | ✔ Yes | 172518.76 |
| exp. var., out | absInStrT_sqrt | 0.23610694 | 0.0014788851 | 0.000000e+00 | ✔ Yes | 172518.76 |
| exp. var., out | absOutStrT_sqrt | -0.07599671 | 0.0014553914 | 0.000000e+00 | ✔ Yes | 172518.76 |
| exp. var., in + out | (Intercept) | 2.05479175 | 0.0028207627 | 0.000000e+00 | ✔ Yes | 144773.85 |
| exp. var., in + out | absInStrT_sqrt | 0.26581550 | 0.0013998517 | 0.000000e+00 | ✔ Yes | 144773.85 |
| exp. var., in + out | absOutStrT_sqrt | -0.04811173 | 0.0013049873 | 3.496844e-296 | ✔ Yes | 144773.85 |

##### 3.2 GLMM: Relative change of expression variance

To investigate whether local network centrality measures affect the evolution of expression variance, we fitted a linear mixed-effects model with the same formula as Eq. 1. with the normalized change of expression variance as the outcome variable. When fitting a model with the assumption of constant variance, the Pearson's residuals were heteroskedastic (Fig S10A). We fitted models with different variance structures and based on Akaike's Information Criterion chose the model with the exponential function of the node abstrength as the variance structure. Pearson's residuals of the chosen and all other fitted models are shown in Fig S10B-E. Changing the variance structure did not change the significance or the effect of the fixed variables (Table S3). The variance inflation factor (VIF), a measure of collinearity of explanatory variables, was 1.06. A VIF value lower than 3 indicates that the statistical significance of the inferred effects is reliable in spite of collinearity.

**Table S3. Different variance structures do not affect the sign of the effect and significance in linear mixed-effects models with relative change of expression variance after selection as a response variable.** The results of models with different variance structures are shown in the table. The effect size differs by a small margin, but the sign and significance remain the same regardless of variance structure. The model with the variance structure as an exponential function of instrength had the lowest Akaike's Information Criterion and was chosen as the best model. Abbreviations: const. var. - constant variance; power var., in + out; variance as a power function of instrength; exp. var., in - variance as an exponential function of instrength; exp. var., out - variance as an exponential function of outstrength; exp. var., in + out - variance as an exponential function of instrength and outstrength; absInStrT_sqrt - absolute instrength, square-root transformed; absOutStrT_sqrt - absolute outstrength, square-root transformed.

| Model | Predictors | Value | Std.Error | DF | t.value | p.value | AIC | p.significant |
|---|---|---|---|---|---|---|---|---|
| const. var. | (Intercept) | 0.337916277 | 0.0013430352 | 72373 | 251.60641 | 0.000000e+00 | -144342.1 | ✔ Yes |
| const. var. | absInStrT_sqrt | -0.008901309 | 0.0004334489 | 72373 | -20.53600 | 1.898873e-93 | -144342.1 | ✔ Yes |
| const. var. | absOutStrT_sqrt | -0.039686466 | 0.0004269809 | 72373 | -92.94669 | 0.000000e+00 | -144342.1 | ✔ Yes |
| power var., in + out | (Intercept) | 0.353325888 | 0.0011743341 | 72373 | 300.87339 | 0.000000e+00 | -149555.2 | ✔ Yes |
| power var., in + out | absInStrT_sqrt | -0.013152971 | 0.0004173796 | 72373 | -31.51321 | 1.686444e-216 | -149555.2 | ✔ Yes |
| power var., in + out | absOutStrT_sqrt | -0.047848893 | 0.0003958800 | 72373 | -120.86718 | 0.000000e+00 | -149555.2 | ✔ Yes |
| ★ exp. var., in | (Intercept) | 0.339920854 | 0.0011538478 | 72373 | 294.59766 | 0.000000e+00 | -153714.6 | ✔ Yes |
| ★ exp. var., in | absInStrT_sqrt | -0.002698555 | 0.0004280085 | 72373 | -6.30491 | 2.900180e-10 | -153714.6 | ✔ Yes |
| ★ exp. var., in | absOutStrT_sqrt | -0.046068532 | 0.0003902311 | 72373 | -118.05450 | 0.000000e+00 | -153714.6 | ✔ Yes |
| exp. var., out | (Intercept) | 0.338768771 | 0.0013394715 | 72373 | 252.91227 | 0.000000e+00 | -144376.3 | ✔ Yes |
| exp. var., out | absInStrT_sqrt | -0.009392597 | 0.0004342054 | 72373 | -21.63169 | 1.926129e-103 | -144376.3 | ✔ Yes |
| exp. var., out | absOutStrT_sqrt | -0.039910911 | 0.0004274997 | 72373 | -93.35893 | 0.000000e+00 | -144376.3 | ✔ Yes |
| exp. var., in + out | (Intercept) | 0.348060641 | 0.0012043388 | 72373 | 289.00558 | 0.000000e+00 | -150295.3 | ✔ Yes |
| exp. var., in + out | absInStrT_sqrt | -0.011395088 | 0.0004347183 | 72373 | -26.21258 | 9.703240e-151 | -150295.3 | ✔ Yes |
| exp. var., in + out | absOutStrT_sqrt | -0.046273751 | 0.0004127882 | 72373 | -112.10047 | 0.000000e+00 | -150295.3 | ✔ Yes |

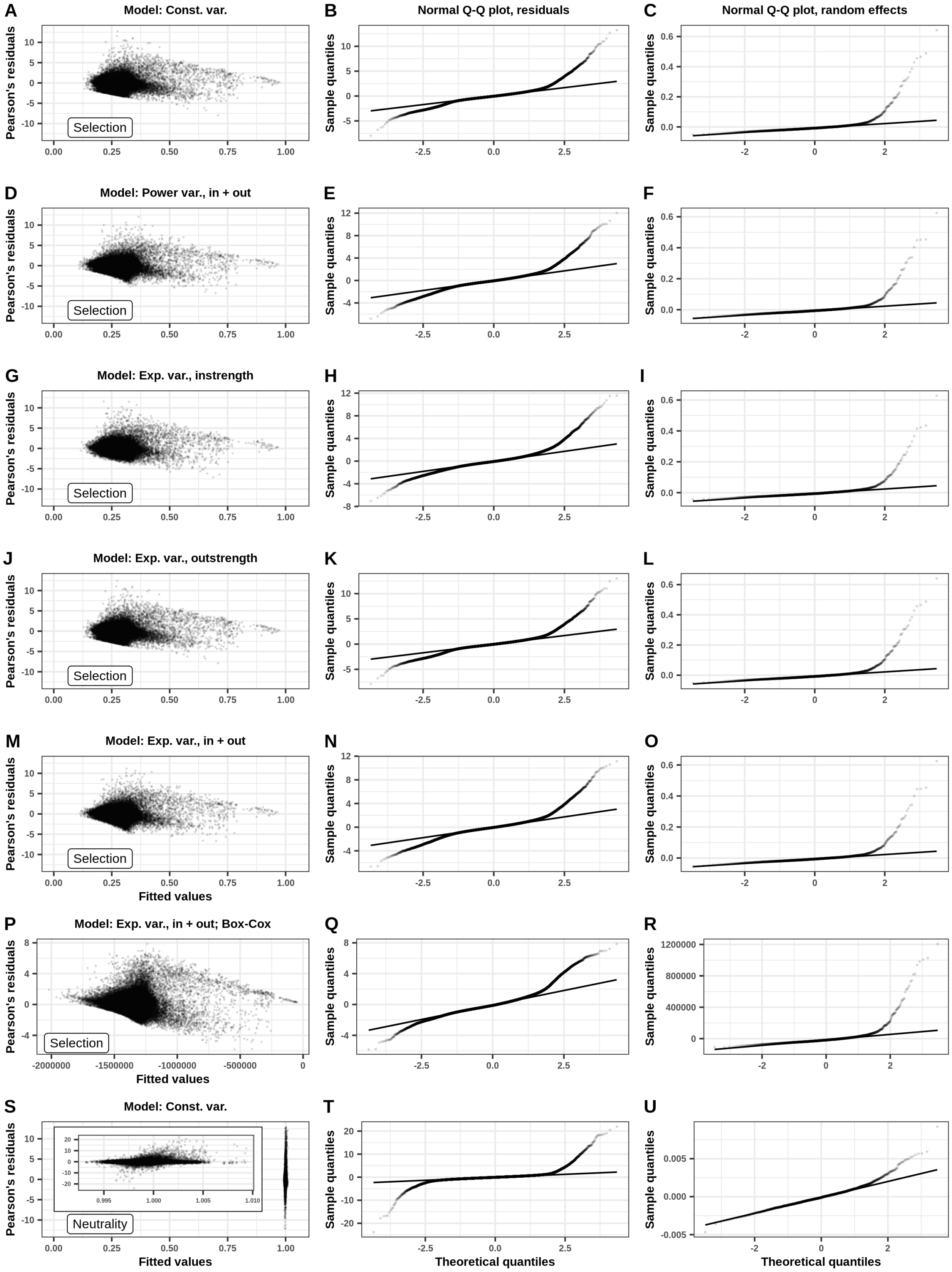

**Fig S10. Diagnostics of linear mixed-effects models with relative change of expression variance after selection as a response variable and different variance structures.**

**Fig S10.** Continued. **A-C** Plot of Pearson's residuals vs. fitted values (A), Q-Q plot of Pearson's residuals (B), Q-Q plot of random effects (C) of a model with no variance structure. **D-F** Pearson's residuals vs. fitted values (D), Q-Q plot of standardized Pearson's residuals (E), Q-Q plot of random effects (C) of a model with a variance structure modelled as a power function of instrength and outstrength. **G-I** Pearson's residuals vs. fitted values (G), Q-Q plot of standardized Pearson's residuals (H), Q-Q plot of random effects (I) of a model with a variance structure modelled as an exponential function of instrength. **J-L** Pearson's residuals vs. fitted values (J), Q-Q plot of standardized Pearson's residuals (K), Q-Q plot of random effects (L) of a model with a variance structure modelled as an exponential function of outstrength. **M-O** Pearson's residuals vs. fitted values (M), Q-Q plot of standardized Pearson's residuals (N), Q-Q plot of random effects (O) of a model with a variance structure modelled as an exponential function of instrength and outstrength. **P-R** Pearson's residuals vs. fitted values (P), Q-Q plot of standardized Pearson's residuals (Q), Q-Q plot of random effects (R) of a model with a variance structure modelled as an exponential function of instrength and outstrength and with the explanatory variables transformed with the Box-Cox transform. **S-U** Pearson's residuals vs. fitted values (M), Q-Q plot of standardized Pearson's residuals (N), Q-Q plot of random effects (O) of a model with constant variance structure fitted on the dataset of populations evolved under neutrality.

##### 3.3 GLMM: Selective pressure

To investigate whether local network centrality measures affect the strength of selective pressure acting on genes, we fitted a linear mixed-effects model with the same formula as Eq. 1. with the selective pressure as the outcome variable. When fitting a model with the assumption of constant variance, the Pearson's residuals were heteroskedastic (Fig S11A). We fitted models with different variance structures and based on Akaike's Information Criterion chose the model with the exponential function of the node abstrength as the variance structure. Pearson's residuals of the chosen and all other fitted models are shown in Fig S11B-E. Changing the variance structure did not change the significance or the effect of the fixed variables (Table S4). The variance inflation factor (VIF), a measure of collinearity of explanatory variables, was 1.03. A VIF value lower than 3 indicates that the statistical significance of the inferred effects is reliable in spite of collinearity.

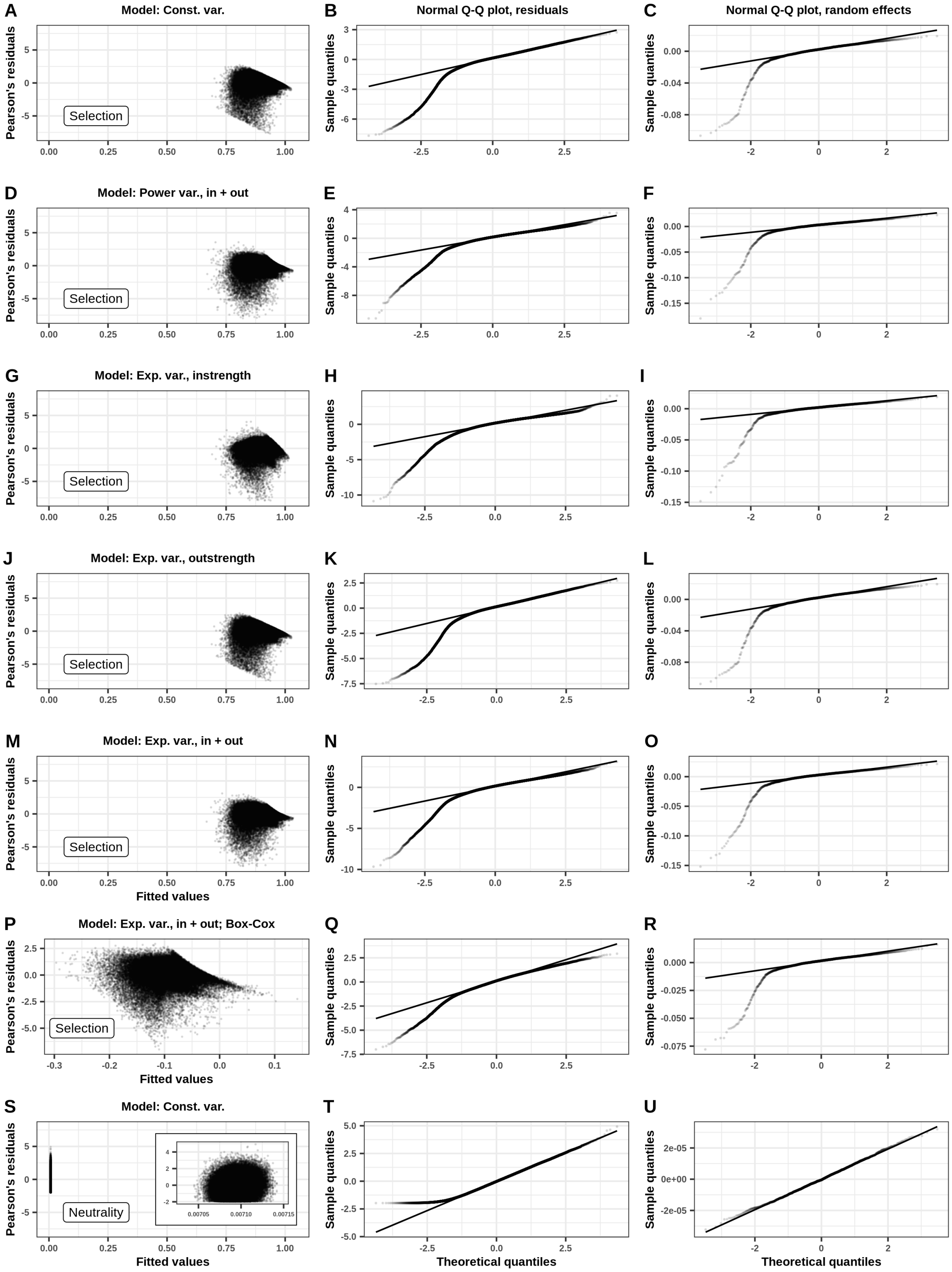

**Fig S11. Diagnostics of linear mixed-effects models with selective pressure as a response variable and different variance structures.**

**Fig S11.** Continued. **A-C** Plot of Pearson's residuals vs. fitted values (A), Q-Q plot of Pearson's residuals (B), Q-Q plot of random effects (C) of a model with no variance structure. **D-F** Pearson's residuals vs. fitted values (D), Q-Q plot of standardized Pearson's residuals (E), Q-Q plot of random effects (C) of a model with a variance structure modelled as a power function of instrength and outstrength. **G-I** Pearson's residuals vs. fitted values (G), Q-Q plot of standardized Pearson's residuals (H), Q-Q plot of random effects (I) of a model with a variance structure modelled as an exponential function of instrength. **J-L** Pearson's residuals vs. fitted values (J), Q-Q plot of standardized Pearson's residuals (K), Q-Q plot of random effects (L) of a model with a variance structure modelled as an exponential function of outstrength. **M-O** Pearson's residuals vs. fitted values (M), Q-Q plot of standardized Pearson's residuals (N), Q-Q plot of random effects (O) of a model with a variance structure modelled as an exponential function of instrength and outstrength. **P-R** Pearson's residuals vs. fitted values (P), Q-Q plot of standardized Pearson's residuals (Q), Q-Q plot of random effects (R) of a model with a variance structure modelled as an exponential function of instrength and outstrength and with the explanatory variables transformed with the Box-Cox transform. **S-U** Pearson's residuals vs. fitted values (M), Q-Q plot of standardized Pearson's residuals (N), Q-Q plot of random effects (O) of a model with constant variance structure fitted on the dataset of populations evolved under neutrality.

**Table S4. Different variance structures do not affect the sign of the effect and significance in linear mixed-effects models with selective pressure as a response variable.** The results of models with different variance structures are shown in the table. The effect size differs by a small margin, but the sign and significance remain the same regardless of variance structure. The model with the variance structure as an exponential function of instrength had the lowest Akaike's Information Criterion and was chosen as the best model. Abbreviations: const. var. - constant variance; power var., in + out; variance as a power function of instrength; exp. var., in - variance as an exponential function of instrength; exp. var., out - variance as an exponential function of outstrength; exp. var., in + out - variance as an exponential function of instrength and outstrength; absInStrT_sqrt - absolute instrength, square-root transformed; absOutStrT_sqrt - absolute outstrength, square-root transformed.

| Model | Predictors | Value | Std.Error | DF | t.value | p.value | p.significant | AIC |
|---|---|---|---|---|---|---|---|---|
| const. var. | (Intercept) | 0.88415564 | 0.0006748052 | 63167 | 1310.2382 | 0 | ✔ Yes | -189809.3 |
| const. var. | absInStrT_sqrt | -0.04096498 | 0.0003106663 | 63167 | -131.8617 | 0 | ✔ Yes | -189809.3 |
| const. var. | absOutStrT_sqrt | 0.03588198 | 0.0002908981 | 63167 | 123.3490 | 0 | ✔ Yes | -189809.3 |
| power var., in + out | (Intercept) | 0.87597512 | 0.0005514233 | 63167 | 1588.5710 | 0 | ✔ Yes | -199304.3 |
| power var., in + out | absInStrT_sqrt | -0.03786164 | 0.0002801003 | 63167 | -135.1717 | 0 | ✔ Yes | -199304.3 |
| power var., in + out | absOutStrT_sqrt | 0.03995255 | 0.0002456307 | 63167 | 162.6529 | 0 | ✔ Yes | -199304.3 |
| ★ exp. var., in | (Intercept) | 0.88278537 | 0.0005377403 | 63167 | 1641.6576 | 0 | ✔ Yes | -207009.0 |
| ★ exp. var., in | absInStrT_sqrt | -0.03708753 | 0.0002878319 | 63167 | -128.8513 | 0 | ✔ Yes | -207009.0 |
| ★ exp. var., in | absOutStrT_sqrt | 0.03434249 | 0.0002328101 | 63167 | 147.5129 | 0 | ✔ Yes | -207009.0 |
| exp. var., out | (Intercept) | 0.88370457 | 0.0006732199 | 63167 | 1312.6537 | 0 | ✔ Yes | -189839.4 |
| exp. var., out | absInStrT_sqrt | -0.04101737 | 0.0003111198 | 63167 | -131.8379 | 0 | ✔ Yes | -189839.4 |
| exp. var., out | absOutStrT_sqrt | 0.03629719 | 0.0002910387 | 63167 | 124.7160 | 0 | ✔ Yes | -189839.4 |
| exp. var., in + out | (Intercept) | 0.87761056 | 0.0005922562 | 63167 | 1481.8090 | 0 | ✔ Yes | -199570.5 |
| exp. var., in + out | absInStrT_sqrt | -0.03864219 | 0.0002999411 | 63167 | -128.8326 | 0 | ✔ Yes | -199570.5 |
| exp. var., in + out | absOutStrT_sqrt | 0.03974518 | 0.0002648410 | 63167 | 150.0719 | 0 | ✔ Yes | -199570.5 |

##### 3.4 GLM: Average selective pressure

To investigate whether global network centrality measures affect the average selective pressure acting on genes in networks, we fitted a linear model with the average selective pressure per network as the outcome variable, and the first two principal components as explanatory variables, after performing a principal component analysis on 12 graph-level centrality metrics. Diagnostical plots are shown in Fig S12.

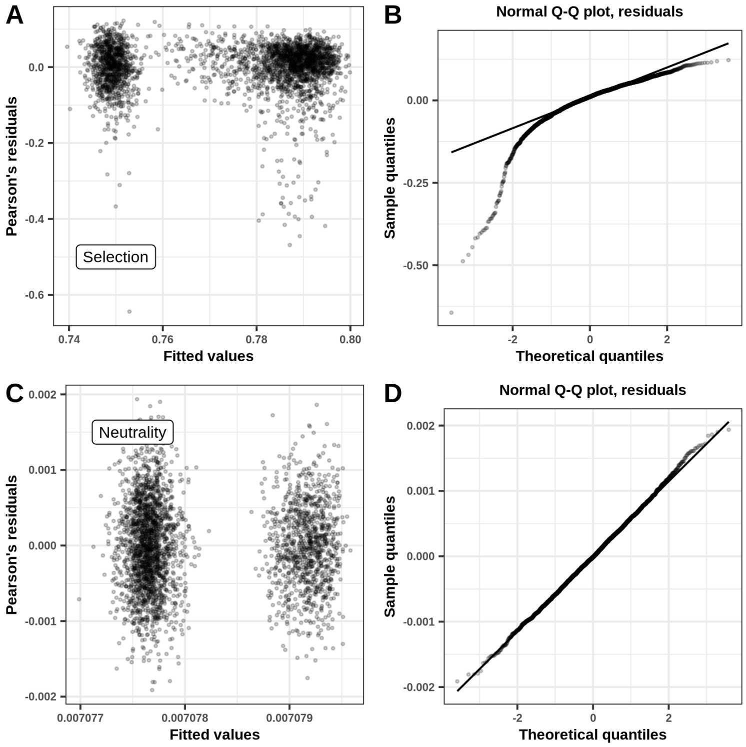

**Fig S12. Diagnostics of linear model with average selective pressure per network as a response variable. A, B** - Pearson's residuals vs. fitted values (A) and Q-Q plot of standardized Pearson's residuals (B) in the model fitted on selected populations. **C, D** - Pearson's residuals vs. fitted values (C) and Q-Q plot of standardized Pearson's residuals (D) in the model fitted on neutral populations.

### 4 Robustness of results in different topology structures

The analysis of the effects of local network metrics on the evolution of expression noise was performed on a dataset of 2,000 random (Erdős–Rényi) network topologies. To check whether our results hold for other network topology types, we performed the same analysis of two additional datasets: 1,000 scale-free (Barabási–Albert) networks, and 1,000 small-world (Watts–Strogatz model) networks. The results of all generalized linear mixed-effects models and mutual information tests are consistent and summarized in Table S5.

**Table S5. The effects and significance of local network centrality metrics are consistent across different topological structures.** The effect size differs by a small margin, but the sign and significance remain the same across different topological structures. Dataset consists of 113,274 genes from 3,000 network topologies.

| Response | Topology | Expl. var. | Beta | p-value (GLMM)[1] | MI | p-value (MI)[2] |
|---|---|---|---|---|---|---|
| Expression variance | ER | Instrength | 0.28 | $< 2.2 \times 10^{-16}$ *** | 0.67 | $10^{-4}$ *** |
| | | Outstrength | -0.02 | $< 2.2 \times 10^{-16}$ *** | 0.05 | $10^{-4}$ *** |
| | BA | Instrength | 0.21 | $< 2.2 \times 10^{-16}$ *** | 0.34 | $10^{-4}$ *** |
| | | Outstrength | -0.06 | $< 2.2 \times 10^{-16}$ *** | 0.11 | $10^{-4}$ *** |
| | WS | Instrength | 0.25 | $< 2.2 \times 10^{-16}$ *** | 0.43 | $10^{-4}$ *** |
| | | Outstrength | -0.05 | $< 2.2 \times 10^{-16}$ *** | 0.05 | $10^{-4}$ *** |
| Rel. change of expr. variance | ER | Instrength | -0.003 | $2.9 \times 10^{-10}$ *** | 0.09 | $10^{-4}$ *** |
| | | Outstrength | -0.046 | $< 2.2 \times 10^{-16}$ *** | 0.14 | $10^{-4}$ *** |
| | BA | Instrength | -0.041 | $< 2.2 \times 10^{-16}$ *** | 0.19 | $10^{-4}$ *** |
| | | Outstrength | -0.027 | $< 2.2 \times 10^{-16}$ *** | 0.26 | $10^{-4}$ *** |
| | WS | Instrength | 0.004 | $< 2.6 \times 10^{-7}$ *** | 0.09 | $10^{-4}$ *** |
| | | Outstrength | -0.039 | $< 2.2 \times 10^{-16}$ *** | 0.08 | $10^{-4}$ *** |
| Probability of responding to selection | ER | Instrength | -1.87 | $< 2.2 \times 10^{-16}$ *** | — | — |
| | | Outstrength | -0.08 | $< 2.2 \times 10^{-16}$ *** | — | — |
| | BA | Instrength | -2.01 | $< 2.2 \times 10^{-16}$ *** | — | — |
| | | Outstrength | -0.4 | $< 2.2 \times 10^{-16}$ *** | — | — |
| | WS | Instrength | -1.88 | $< 2.2 \times 10^{-16}$ *** | — | — |
| | | Outstrength | -0.16 | $3.9 \times 10^{-11}$ *** | — | — |
| Gene-specific selective pressure | ER | Instrength | -0.04 | $< 2.2 \times 10^{-16}$ *** | 0.19 | $10^{-4}$ *** |
| | | Outstrength | 0.03 | $< 2.2 \times 10^{-16}$ *** | 0.31 | $10^{-4}$ *** |
| | BA | Instrength | -0.02 | $< 2.2 \times 10^{-16}$ *** | 0.14 | $10^{-4}$ *** |
| | | Outstrength | 0.02 | $< 2.2 \times 10^{-16}$ *** | 0.54 | $10^{-4}$ *** |
| | WS | Instrength | -0.04 | $< 2.2 \times 10^{-16}$ *** | 0.17 | $10^{-4}$ *** |
| | | Outstrength | 0.03 | $< 2.2 \times 10^{-16}$ *** | 0.2 | $10^{-4}$ *** |

[1] Coefficients and their significance were computed using linear mixed-effects model (see Methods).
[2] Mutual information p-values were computed using a Monte Carlo permutation test with 10,000 permutations. Asterisks indicate statistical significance: n.s. - p-value > 0.05; * - p-value ≤ 0.05; ** - p-value ≤ 0.01; *** - p-value ≤ 0.001; **** - p-value ≤ 0.0001.

#### 5 Filtered datasets

We performed the analyses on two additional datasets to get a clearer picture of the effects of instrength and outstrength on the expression noise metrics. The first filtered dataset is a dataset in which genes that are both regulators and regulated were removed, *i.e.* it consists exclusively of pure regulators (genes that regulate others and are not being regulated) and purely regulated genes (genes that are being regulated and do not regulate other genes). The second filtered dataset is a dataset that consists exclusively of genes that are both regulators and regulated, *i.e.* in this dataset pure regulators and purely regulated genes have been removed. The effects and significance of the two local centrality metrics are consistent in analyses of expression variance, relative change of expression variance and gene-specific selective pressure (Table S6). In the unfiltered and second filtered dataset we found a significant small negative effect of outstrength on the probability of responding to selection. However, the negative effect is lost when we analysed only the genes that are either regulators or regulated genes (Filtered 1 dataset), and we observed a significant strong positive effect of outstrength on the probability of responding to selection. We concluded that instrength in genes that are both regulators and regulated influences the effect of outstrength on the probability of responding to selection and that there are complex interactions between the two centrality metrics. However, when there are no genes that have both instrength and outstrength the effects are clear and instrength has a strongly negative effect, while outstrength has a strongly positive effect on the probability of a gene to respond to selection.

**Table S6. Filtered and unfiltered datasets.** Filtered 1 dataset is dataset in which genes that are both regulators and regulated were removed, *i.e.* it consists exclusively of pure regulators (genes that regulate others and are not being regulated) and purely regulated genes (genes that are being regulated and do not regulate other genes). Filtered 1 dataset consists of 43,214 genes from 2,000 random network topologes. Unfiltered dataset consists of 148,886 genes from 2,000 random network topologes. Filtered 2 dataset consists exclusively of genes that are both regulators and regulated, *i.e.* in this dataset pure regulators and purely regulated genes have been removed. Filtered 2 dataset consists of 105,672 genes from 2,000 random network topologes.

| **Response** | **Dataset** | **Expl. var.** | **Beta** | **p-value (GLMM)**[1] | **MI** | **p-value (MI)**[2] |
|---|---|---|---|---|---|---|
| Expression variance | Filtered 1 | Instrength | 0.29 | $< 2.2 \times 10^{-16}$ *** | 0.8 | $10^{-4}$ *** |
| | | Outstrength | $7.8 \times 10^{-4}$ | $< 2.6 \times 10^{-9}$ *** | 0.61 | $10^{-4}$ *** |
| | Unfiltered | Instrength | 0.28 | $< 2.2 \times 10^{-16}$ *** | 0.67 | $10^{-4}$ *** |
| | | Outstrength | -0.022 | $< 2.2 \times 10^{-16}$ *** | 0.05 | $10^{-4}$ *** |
| | Filtered 2 | Instrength | 0.24 | $< 2.2 \times 10^{-16}$ *** | 0.43 | $10^{-4}$ *** |
| | | Outstrength | -0.08 | $< 2.2 \times 10^{-16}$ *** | 0.02 | $10^{-4}$ *** |
| Rel. change of expr. variance | Filtered 1 | Instrength | -0.033 | $< 2.2 \times 10^{-16}$ *** | 0.29 | $10^{-4}$ *** |
| | | Outstrength | -0.073 | $< 2.2 \times 10^{-16}$ *** | 0.34 | $10^{-4}$ *** |
| | Unfiltered | Instrength | -0.003 | $2.9 \times 10^{-10}$ *** | 0.09 | $10^{-4}$ *** |
| | | Outstrength | -0.046 | $< 2.2 \times 10^{-16}$ *** | 0.14 | $10^{-4}$ *** |
| | Filtered 2 | Instrength | -0.017 | $< 2.2 \times 10^{-16}$ *** | 0.07 | $10^{-4}$ *** |
| | | Outstrength | -0.035 | $< 2.2 \times 10^{-16}$ *** | 0.08 | $10^{-4}$ *** |
| Probability of responding to selection | Filtered 1 | Instrength | -1.94 | $< 2.2 \times 10^{-16}$ *** | — | — |
| | | Outstrength | 1.55 | $< 9.79 \times 10^{-11}$ *** | — | — |
| | Unfiltered | Instrength | -1.87 | $< 2.2 \times 10^{-16}$ *** | — | — |
| | | Outstrength | -0.08 | $< 2.2 \times 10^{-16}$ *** | — | — |
| | Filtered 2 | Instrength | -1.79 | $< 2.2 \times 10^{-16}$ *** | — | — |
| | | Outstrength | -0.25 | $< 2.2 \times 10^{-16}$ *** | — | — |
| Gene-specific selective pressure | Filtered 1 | Instrength | -0.05 | $< 2.2 \times 10^{-16}$ *** | 0.63 | $10^{-4}$ *** |
| | | Outstrength | 0.03 | $< 2.2 \times 10^{-16}$ *** | 0.72 | $10^{-4}$ *** |
| | Unfiltered | Instrength | -0.04 | $< 2.2 \times 10^{-16}$ *** | 0.1 | $10^{-4}$ *** |
| | | Outstrength | 0.03 | $< 2.2 \times 10^{-16}$ *** | 0.05 | $10^{-4}$ *** |
| | Filtered 2 | Instrength | -0.04 | $< 2.2 \times 10^{-16}$ *** | 0.1 | $10^{-4}$ *** |
| | | Outstrength | 0.03 | $< 2.2 \times 10^{-16}$ *** | 0.17 | $10^{-4}$ *** |

[1] Coefficients and their significance were computed using linear mixed-effects model (see Methods).
[2] Mutual information p-values were computed using a Monte Carlo permutation test with 10,000 permutations. Asterisks indicate statistical significance: n.s. - p-value > 0.05; * - p-value ≤ 0.05; ** - p-value ≤ 0.01; *** - p-value ≤ 0.001; **** - p-value ≤ 0.0001.

#### References

1. James G, Witten D, Hastie T, Tibshirani R. An Introduction to Statistical Learning: with Applications in R. Springer Texts in Statistics. Springer US;. Available from: https://link.springer.com/10.1007/978-1-0716-1418-1.
